# Representation Learning to Effectively Integrate and Interpret Omics Data

**DOI:** 10.1101/2023.04.23.537975

**Authors:** Sara Masarone

**Affiliations:** The Alan Turing Institute, Queen Mary University of London, London, UK

## Abstract

The last decade has seen an increase in the amount of high throughput data available to researchers. While this has allowed scientists to explore various hypotheses and research questions, it has also highlighted the importance of data integration to facilitate knowledge extraction and discovery. Although many strategies have been developed over the last few years, integrating data whilst generating an interpretable embedding still remains challenging due to difficulty in regularisation, especially when using deep generative models. Thus, we introduce a framework called Regularised Multi-View Variational Autoencoder (RMV-VAE) to integrate different omics data types whilst allowing researchers to obtain more biologically meaningful embeddings.

## 1 Introduction

In the last few decades, technological progress has yielded a large quantity of heterogeneous high-throughput data which has provided information about processes underpinning various diseases of interest. This has allowed researchers to unveil molecular patterns in genomics, proteomics and lipidomics leading to a much more detailed understanding of the pathways and systems that regulate the immune response in humans. The sudden increase in data highlighted, at the same time, the importance of integrating different types of molecular data together to enhance pattern discovery and improve patient stratification.

Due to economic limitations, studies have historically focused on the use of unimodal data, thus concentrating on a single aspect of the Dna-Rna-Protein paradigm. While this approach has proven valuable to gain insights into various conditions, the lack of robust data integration has limited discoveries by failing to analyse biological systems as a whole. This has been a limiting factor since, when looking at a particular data type, we are just analysing a snapshot of the condition of interest and, therefore, it can only provide us with limited insights on the complex processes underpinning a disease. Being mostly unfeasible to extensively monitor omics changes over time, we can resort to the use of multiple data types to get a more complete picture of the pathways being modulated in the process [1].

In this paper, we consider different omics data types as different “views” of the same system [2, 3]. We aim to use a fusion approach to integrate these views together whilst obtaining an interpretable low dimensional data representation, or embedding, where points close in space share similar characteristics. This is crucial when trying to use model’s embeddings for downstream tasks such as clustering or when we aim to obtain clinically meaningful groups.

In the following sections we will explore related work and we will present a few experiments that support the use of our framework called Regularised Multi-View Variational Autoencoder (RMV-VAE) on omics datasets. Our approach is summarised in figure 1.

**Figure 1:**
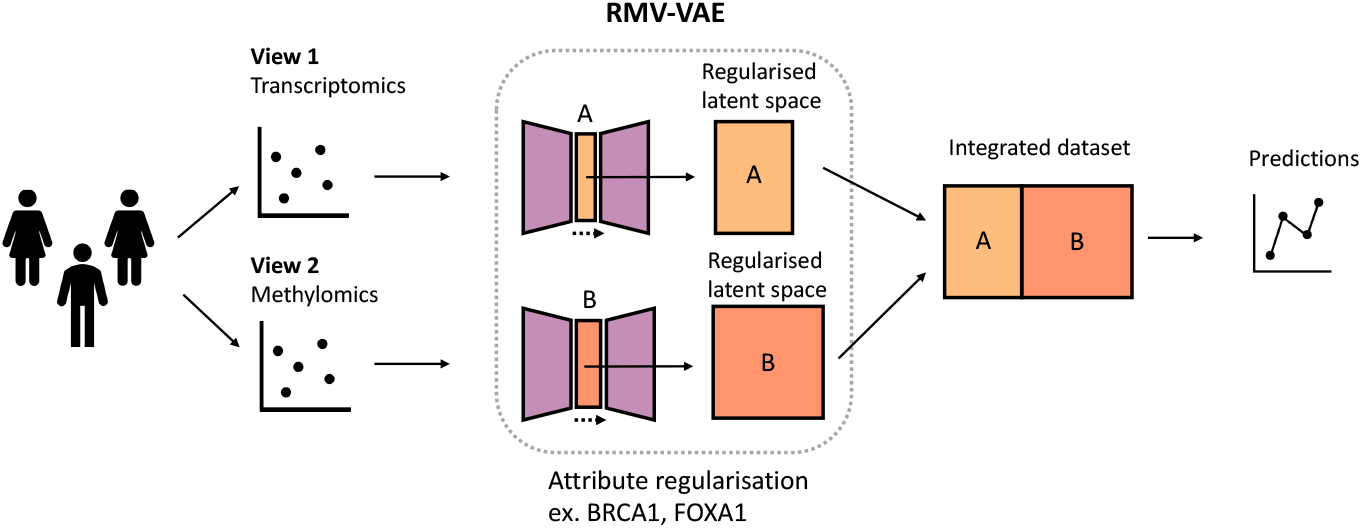
Overview of the RMV-VAE framework. The framework is composed of two Variational Autoencoders that take datasets as input and generate a regularised low dimensional representation of the data. The regularisation is achieved by adding an attribute specific loss to the standard VAE loss. The embeddings are then merged and subsequently used to predict outcomes.

### 1.1 Related works

Data integration and dimensionality reduction are central aspects of many data analysis pipelines both in bioinformatics and other fields [4]. Among many models, Variational Autoencoders (VAEs) have been shown to be useful at solving a variety of problems such as data compression, de-noising, learning transferable representations of data and data fusion [5, 6, 7]. Nevertheless, a great limitation of using VAEs in biology has been the inability to obtain regularised, and hence interpretable, embeddings where points close in the original feature space are also close in the latent space, limiting their use in clinical practice [8, 9].

In the context of omics data integration, OmiEmbed and SubOmiEmbed have shown the potential of representation learning and self-representation learning at integrating data and performing downstream predictions in a unified framework [10, 11]. Other implementations, such as Multi-Encoder VAE applied to single cell data, have highlighted the importance of feeding different views of the same data to obtain more robust data representations and more separable classes [3].

In single cells RNA-Seq experiments, Graph Neural Networks (GNN) such as GLUE (graph-linked unified embedding) have also shown their potential at integrating datasets together by modeling regulatory interactions across omics layers explicitly [12]. Other models such as scVI have also been developed showing competitive results on single cell data [13]. While we acknowledge the performance and usefulness of GNNs in single cell RNA-Seq data, we recognise that adapting these models to bulk RNA-Seq data (used in these experiments) is a non-trivial task as it requires knowledge of the interaction between the different data-types to allow the construction of a meaningful graph.

Thus, our work takes inspiration from the aforementioned VAE research, expanding it on the generation of an interpretable, regularisable latent space. In particular, we designed this model to allow researchers to not only perform downstream predictions, but also regularise the embeddings by a gene or protein known to be important in that condition, thus improving patient stratification and treatment.

## 2 Proposed method

We propose a probabilistic framework that can be extended to *N* datasets allowing for dynamic integration of omics data. The framework comprises two VAEs each composed of an encoder and a decoder. Each VAE takes a dataset as input (*X*) and generates a low dimensional, or latent, representation of the data (*z*). Once the data has been compressed, it is reconstructed by passing it through a decoder which is symmetrical to the encoder. VAEs work by minimising the evidence lower bound (ELBO). By using a reconstruction loss between the model’s input *X* and the output 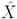 and a KL divergence between the encoded data and a Normal distribution, the model is forced to learn the “real” signal present in the data, thus prioritising signal to noise 2. In addition, the KL divergence forces the latent space to be more regularised than it would be just by using similar compression methods such as Autoencoders.

To encode an attribute *a* along one dimension *r* of the latent space (*z*) we used a regularisation loss introduced by Pati et al. [14] where, as we traverse along *r*, the value of *a* increases. The regularisation is achieved by adding an additional term to the loss of this model. This allows users to generate embeddings that “order” patients by the expression of a specific variable of interest (eg. BRCA1 gene for breast cancer) improving results’ interpretation.

The attribute loss is formulated as follows:

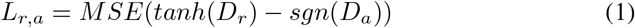

where *MSE* is the Mean Squared Error, *D*_*r*_ is the distance matrix computed for the regularised dimension *r*, and *D*_*a*_ is the distance matrix computed for the attribute we wish to encode along a dimension of *z. Tanh* and *sgn* are the hyperbolic tangent function and sign function respectively. This additional term is added to the standard Reconstruction and KL Loss 2 and forces a monotonic relationship between the encoded attribute and a dimension of *z* allowing users to obtain a more meaningful^1^ and structured latent space.

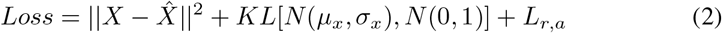

### 2.1 Training overview

The models used in training were built using Tensorflow v. 2.9.2. To ensure the generation of interpretable embeddings, we tested the size of the networks, as well as the hidden layers and the batch size manually, as in this particular setting optimising for performance would not necessarily imply an increase in the embedding interpretability, a problem that has been previously discussed in the literature [15].

We finally tested a range of nodes for the hidden layers [100, 80, 60, 40, 20, 10] to find the most suitable architecture. Model training was performed on CPUs. Average run time for the BRCA dataset (VAE/RMV-VAE) was 30s and 28s respectively, while for the PAAD dataset was 11.8s (VAE) and 28.06s (RMV-VAE) respectively. You can find our github repository here.

## 3 Experiments and results

We tested our framework on two diseases, breast cancer and pancreatic cancer. Both experiments were performed using transcriptomics and methylomics datasets. We assessed the ability of our framework to integrate different views of the same system ^2^ and then explored the interpretability of the latent space generated (see appendix for additional experiments).

### 3.1 Experiment 1 - breast cancer datasets

We retrieved transcriptomics and methylomics datasets from the UCSC Xena TCGA data repository containing 1212 and 885 patients respectively [16]. We preprocessed the data by only selecting the patients present in both datasets reaching a total number of 787 patients. We preprocessed counts using EdgeR [17] and then scaled the datasets before model fitting.

We started by looking at survival as outcome of interest. To encode the transcriptomics, we used a two-layers Neural Network (NN) for both the encoder and the decoder while to encode the methylomics we used a three-layers NN. Given the importance of the BRCA1 and TMEM101 genes in breast cancer survival, we chose them to regularise the latent space of each dataset respectively. We then generated the final embedding that we used for downstream predictions. We evaluated the embedding on prediction tasks such as predicting survival and we compared the results to the ones obtained using individual datasets or a standard VAE. As shown in table 1, the integration obtained by RMV-VAE outperformed the standard VAE as well as individual datasets in predicting survival using a Random Forest Classifier with five folds cross validation (mean accuracy 0.87 vs 0.85 e AUC of 0.62 vs 0.56).

**Table 1:** Results: predicting survival and ER status in breast cancer

| Model | Survival (acc.) | ER (acc.) | Survival (AUC) | ER (AUC) |
| --- | --- | --- | --- | --- |
| RMV-VAE transcriptomics (t) | $0.86 \pm 0.01$ | $0.81 \pm 0.06$ | $0.59 \pm 0.10$ | $0.69 \pm 0.04$ |
| RMV-VAE methylomics (m) | $0.82 \pm 0.02$ | $0.84 \pm 0.03$ | $0.54 \pm 0.05$ | $0.88 \pm 0.04$ |
| **RMV-VAE (t+m)** | **$0.87 \pm 0.01$** | **$0.88 \pm 0.02$** | **$0.62 \pm 0.11$** | **$0.91 \pm 0.05$** |
| VAE transcriptomics (t) | $0.84 \pm 0.03$ | $0.71 \pm 0.03$ | $0.55 \pm 0.04$ | $0.66 \pm 0.03$ |
| VAE methylomics (m) | $0.81 \pm 0.01$ | $0.88 \pm 0.02$ | $0.58 \pm 0.04$ | $0.89 \pm 0.03$ |
| **VAE (t+m)** | **$0.85 \pm 0.02$** | **$0.87 \pm 0.02$** | **$0.56 \pm 0.05$** | **$0.89 \pm 0.03$** |

We repeated the same experiment predicting Estrogen Receptor (ER) status. In this case, we regularised the latent space by two genes known to play key roles in breast cancer’s ER status: FOXA1 and AGR3. We then generated the final embedding that was used for predictions. Results showed that our framework outperformed a standard VAE (using 5 folds CV, AUC of 0.91 vs 0.89 and mean accuracy of 0.88 vs 0.87) at generating a more organised, low dimensional data representation, as visible in table (see following section for more details) 1.

A comparison of the integration achieved by our framework and a standard VAE is shown in figure 2. The figure shows that the data compression obtained by RMV-VAE allows to better separate ER-positive from ER-negative patients in all datasets, therefore leading to an improvement in the final embedding generated.

**Figure 2:**
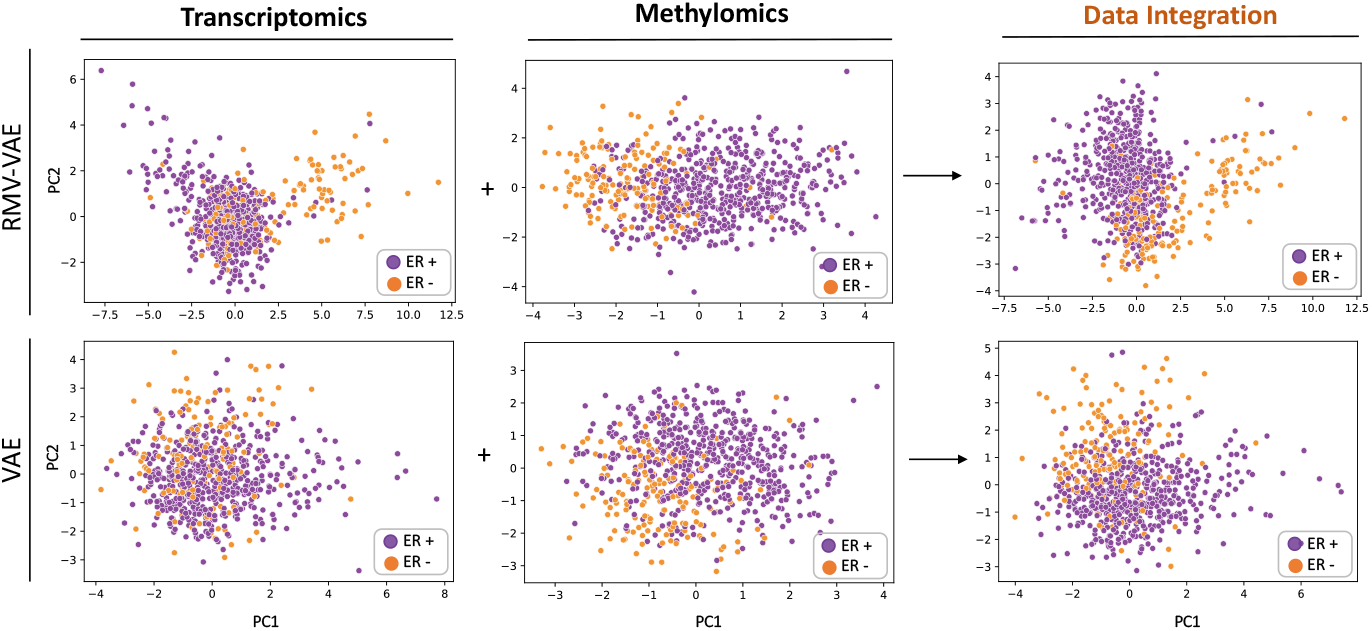
Comparison of the embeddings obtained by RMV-VAE and a standard VAE on BRCA datasets (ER). This figure shows the Principal Component Analysis (PCA) of each embedding (transcriptomics, methylomics and transcriptomics and methylomics integrated). Steps of this pipeline involve: (1) generating an embedding for each dataset first, (2) regularising the embedding by a chosen variable (shown in figure 3) and (3) merging the embeddings to obtain a final representation of both datasets

### 3.2 Experiment 2 - pancreatic cancer datasets

We applied RMV-VAE to the TCGA Pancreatic Adenocarcinoma datasets, consisting of 181 patients. We used a three-layers NN for the transcriptomics and two-layers NN for the methylomics. We preprocessed the datasets by removing methylation sites with low variance and scaled the data before proceeding at generating the individual embeddings. The transcriptomics was preprocessed using standard bioinformatics pipelines (edgeR). To obtain the individual embeddings, we normalised the latent space by KRAS and PTGDR, two driver genes that were shown to have diagnostic and prognostic value in pancreatic cancer [18, 19]. We then combined the embeddings and evaluated performance in predicting survival. As shown in table 2, we obtained a better performance at predicting survival when using this framework compared to a standard VAE (AUC of 0.65 vs 0.57, mean accuracy of 0.62 vs 0.59).

**Table 2:** Results: predicting survival in pancreatic cancer (accuracy and AUC)

| Model | Survival (acc., std) | Survival (AUC, std) |
| --- | --- | --- |
| RMV-VAE counts (c) | $0.6 \pm 0.04$ | $0.65 \pm 0.07$ |
| RMV-VAE methylations (m) | $0.54 \pm 0.06$ | $0.57 \pm 0.10$ |
| **RMV-VAE - (c + m)** | **$0.62 \pm 0.1$** | **$0.65 \pm 0.08$** |
| VAE counts (c) | $0.59 \pm 0.05$ | $0.58 \pm 0.07$ |
| VAE methylations (m) | $0.57 \pm 0.06$ | $0.54 \pm 0.11$ |
| **VAE - (c + m)** | **$0.59 \pm 0.04$** | **$0.57 \pm 0.10$** |

**Table 3:**
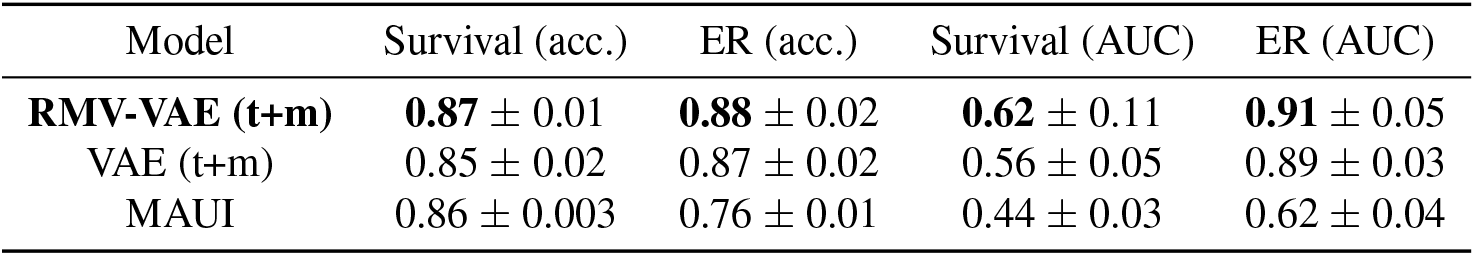
Results: predicting survival and ER status in breast cancer using MAUI

| Model | Survival (acc.) | ER (acc.) | Survival (AUC) | ER (AUC) |
| --- | --- | --- | --- | --- |
| **RMV-VAE (t+m)** | **0.87** ± 0.01 | **0.88** ± 0.02 | **0.62** ± 0.11 | **0.91** ± 0.05 |
| VAE (t+m) | 0.85 ± 0.02 | 0.87 ± 0.02 | 0.56 ± 0.05 | 0.89 ± 0.03 |
| MAUI | 0.86 ± 0.003 | 0.76 ± 0.01 | 0.44 ± 0.03 | 0.62 ± 0.04 |

**Table 4:** Results: predicting survival in pancreatic cancer (accuracy and AUC) using MAUI

| Model | Survival (acc., std) | Survival (AUC, std) |
| --- | --- | --- |
| **RMV-VAE - (t + m)** | **0.62** ± 0.1 | **0.65** ± 0.08 |
| VAE - (t + m) | 0.59 ± 0.04 | 0.57 ± 0.10 |
| MAUI | 0.56 ± 0.08 | 0.51 ± 0.1 |

## 4 Latent space interpretability

One of the key reasons to generate data embeddings using Variational Autoencoders is their ability to compress high dimensional data and obtain a low dimensional representation of it. However, a problem that arises when using VAEs is the difficulty at generating ordered and interpretable latent spaces, as patients that are known to be clinically similar in the original input data are not necessarily close in the VAE embedding, slowing or limiting results’ interpretation. As some genes might be known to play a crucial role in these conditions, we might want to not only compress the data, but also obtain a representation in which patients are ordered by a particular attribute to help relate possible embeddings’ clusters to already established clinical groups.

In the following section we will present some practical examples where this framework might be useful.

### 4.1 Clinical examples

Aggressive phenotypes of ER+ breast cancer are known to be driven by FOXA1 augmentation and expression which leads to activation of key mechanisms that promote metastatic programs [20]. Consequently, it might be useful to consider the expression of this gene when analysing breast cancer. In figure 3, **(A)**, we show the embedding obtained by our model (RMV-VAE) and we compare it to the embedding obtained by a standard VAE and to the original expression data. For each embedding, we plotted the PCA to allow 2d comparisons between them. The panel shows the improvement in class separation (ER+/−) on the top row, and the improvement in regularisation on the bottom row. Results show that our framework allows us to obtain a more structured latent space where patients are ordered by the expression of the FOXA1 gene. This leads to an improvement compared to the embedding of the original data and the standard VAE where patients with high and low expression were not easily separable. Regularising the latent space by FOXA1 we achieve a much clearer separation between ER-negative patients, described in the literature as having low FOXA1 expression levels, and ER-positive patients that tend to display medium-to-high expression of this gene, improving overall data interpretation.

**Figure 3:**
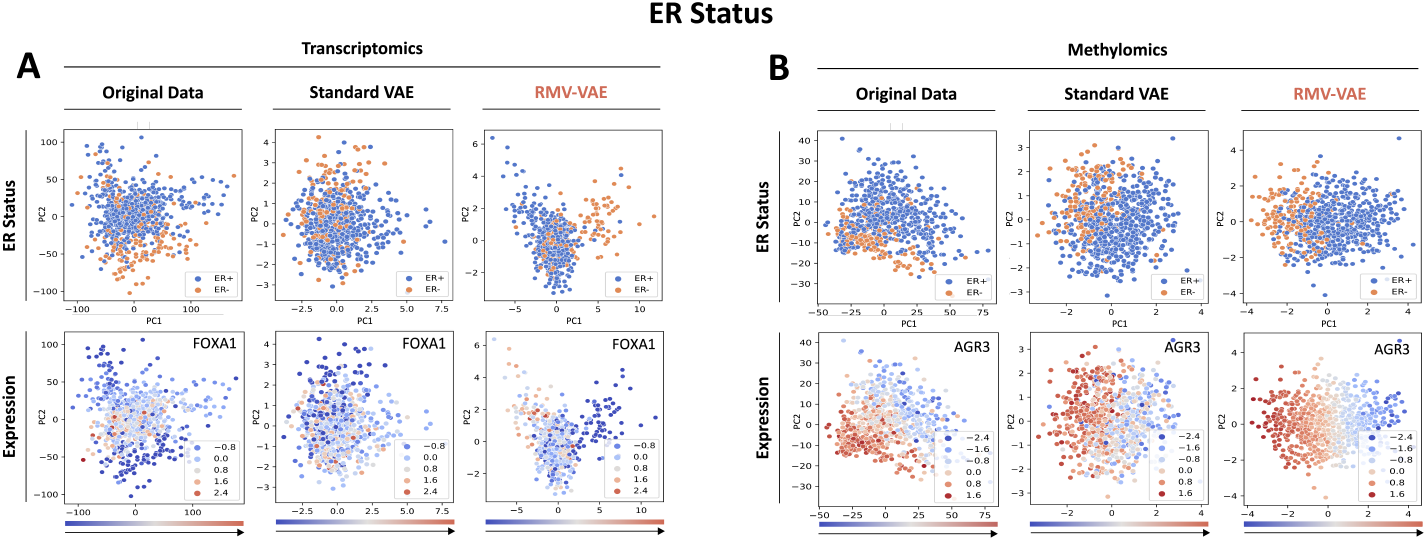
Comparisons between the embedding obtained by our framework (RMVVAE), the original input data and the embedding obtained by a standard VAE. For each embedding, we plotted its PCA to show the results in two dimensions. As shown in the figure, regularising by FOXA1 **(A)** we obtained a better segregation between ER+ and ERin the transcriptomics. In fact, we can now see more clearly that ER- patients have very low levels of FOXA1 compared to ER+ that show, in contrast, mild-to-high levels of this gene. Regularising the methylomics data by AGR3 **(B)** shows an improvement in the separation of the two classes compared to the original data and compared to the classic VAE.

Similarly, the AGR3 gene is a gene known to regulate cell adhesion and migration in breast cancer, two key steps in tumour spreading and metastasis formation [21]. This gene is known to be highly methylated in ER-negative patients and it is therefore a good biomarker to identify these two clinically different groups. Good identification of ER-positive or negative patients would result in more adequate treatment choices and therefore higher chances of survival. In figure 3 section **(B)**, we can compare the embedding obtained by our framework to the original expression data and to the embedding generated by a standard VAE. The figure shows that using our framework we achieved a more organised latent space which results in more separable classes compared to the other embeddings.

The same patterns apply to genes such as BRCA1 and TMEM101 (not shown here). To achieve similar results using a standard VAE, trial and error would be the only option and there would be no guarantee of achieving a good result. Thus, we suggest the use of this framework to allow researchers to have more flexibility in the regularization of their embeddings.

## 5 Discussion

Data integration is a key aspect of biomedical research [1]. However, effective and meaningful data integration is still challenging to achieve due to technical limitations. In this paper, we introduced a new framework, called RMV-VAE, to integrate different data types and obtain more meaningful, low dimensional representations of the data.

We tested our framework on two diseases, breast cancer and pancreatic cancer and we demonstrated the importance of integrating omics data to achieve higher performance at downstream tasks such as predicting ER status or survival.

We compared the results obtained using RMV-VAE to the results obtained using a standard VAE showing an increase in performance when using this framework. In particular, we tested RMV-VAE at predicting survival and ER status for breast cancer, and survival for pancreatic cancer registering an increase in performance in all experiments. The improvement in pancreatic cancer was striking given that this type of cancer is notoriously hard to classify due to the high heterogeneity of the human population.

In addition to integration, which is a key aspect of many data science pipelines, we tried to improve the latent space of VAEs to allow scientists and researchers to use them for downstream tasks. To incorporate their expertise to the embeddings, we formulated an *ah-hoc* regularisation of the latent space to obtain embeddings where patients with similar expression of fundamental genes are found close together. By doing so, we facilitated results interpretation allowing researchers to relate new results to already existent clinical groups.

## 6 Conclusion

Our framework provides a way to integrate and interpret omics data effectively allowing researchers to customise their analyses and explore genes or biomarkers that are known to play central roles in a given disease. In this paper, we introduced this framework, we demonstrated its utility on two real life examples, and we discussed use cases in clinical practice.

## Acknowledgments and Disclosure of Funding

I would like to thank Lorenzo Sani for great discussions. Sara Masarone is supported by The Alan Turing Institute under the EPSRC grant EP/N510129/1.

## A Benchmarking against MAUI

Table of results comparing MAUI (Multi-omics Autoencoder Integration) to RMV-VAE and a standard VAE. MAUI is based on Variational Autoencoders and has been shown to be effective at integrating different omics datasets together [22]. Although we have already benchmarked VAEs before, here we compare it to this specific implementation.

In these experiments, MAUI was set with a hidden layer/latent layer of 1300/600 (survival) and 1400/600 (ER) for the BRCA data and 1300/600 for the PAAD data. The experiments were executed on Google Colab and the run time was on overage 16m.

## Footnotes

1 We intend to obtain a data representation in which patients close in space share similar characteristics such as a similar expression of a gene of interest.

2 Different data modalities.

## References

[1] Xin Luna Dong and Theodoros Rekatsinas. Data Integration and Machine Learning: A Natural Synergy. Proceedings of the 2018 International Conference on Management of Data, 11(12):2094–2097, 2018.

[2] Yingming Li, Ming Yang, and Zhongfei Zhang. A Survey of Multi-View Representation Learning. IEEE Trans Knowl Data Eng, 31(10):1863–1883, 10 2019.

[3] Luke Ternes, Mark Dane, Sean Gross, Marilyne Labrie, Gordon Mills, Joe Gray, Laura Heiser, and Young Hwan Chang. A multi-encoder variational autoencoder controls multiple transformational features in single-cell image analysis. Commun Biol, 5(1):255, 3 2022.

[4] Zhang Zhang, Vladimir B., Jun Yu, Kei-Hoi Cheung, and Jeffrey P. Data integration in bioinformatics: current efforts and challenges. In Mahmood A. Mahdavi, editor, Bioinformatics - Trends and Methodologies. InTech, 11 2011.

[5] Daniel P Gomari, Annalise Schweickart, Leandro Cerchietti, Elisabeth Paietta, Hugo Fernandez, Hassen Al-Amin, Karsten Suhre, and Jan Krumsiek. Variational autoencoders learn transferrable representations of metabolomics data. Commun Biol, 5(1):645, 6 2022.

[6] Diederik P Kingma and Max Welling. An introduction to variational autoencoders. FNT in Machine Learning, 12(4):307–392, 2019.

[7] Sören Richard Stahlschmidt, Benjamin Ulfenborg, and Jane Synnergren. Multimodal deep learning for biomedical data fusion: a review. Brief Bioinformatics, 23(2), 3 2022.

[8] Gaëtan Hadjeres, Frank Nielsen, and François Pachet. GLSR-VAE: Geodesic Latent Space Regularization for Variational AutoEncoder Architectures. arXiv, 2017.

[9] Hao Fu, Chunyuan Li, Xiaodong Liu, Jianfeng Gao, Asli Celikyilmaz, and Lawrence Carin. Cyclical Annealing Schedule: A Simple Approach to Mitigating KL Vanishing. Technical report.

[10] Xiaoyu Zhang, Yuting Xing, Kai Sun, and Yike Guo. OmiEmbed: A Unified Multi-Task Deep Learning Framework for Multi-Omics Data. Cancers (Basel), 13(12), 6 2021.

[11] Sayed Hashim, Muhammad Ali, Karthik Nandakumar, and Mohammad Yaqub. SubOmiEmbed: Self-supervised Representation Learning of Multi-omics Data for Cancer Type Classification. arXiv, 2022.

[12] Zhi-Jie Cao and Ge Gao. Multi-omics single-cell data integration and regulatory inference with graph-linked embedding.

[13] Adam Gayoso, Romain Lopez, Galen Xing, Pierre Boyeau, Valeh Valiollah Pour Amiri, Justin Hong, Katherine Wu, Michael Jayasuriya, Edouard Mehlman, Maxime Langevin, Yining Liu, Jules Samaran, Gabriel Misrachi, Achille Nazaret, Oscar Clivio, Chenling Xu, Tal Ashuach, Mariano Gabitto, Mohammad Lotfol-lahi, Valentine Svensson, Eduardo da Veiga Beltrame, Vitalii Kleshchevnikov, Carlos Talavera-López, Lior Pachter, Fabian J. Theis, Aaron Streets, Michael I. Jordan, Jeffrey Regier, and Nir Yosef. A Python library for probabilistic analysis of single-cell omics data. Nature Biotechnology 2022 40:2, 40(2):163–166, 2 2022.

[14] Ashis Pati and Alexander Lerch. Latent Space Regularization for Explicit Control of Musical Attributes. Proc Int Conf Mach Learn, 2019.

[15] Jonathan Chang, Jordan Boyd-Graber, Sean Gerrish, Chong Wang, and David M Blei. Reading Tea Leaves: How Humans Interpret Topic Models. Advances in Neural Information Processing Systems, 22, 2009.

[16] Goldman M., Craft B., Hastie Mim, H Caicedo H., Daniel A Hashimoto, Julio C Caicedo, Alex Pentland, and Gary P Pisano. Visualizing and interpreting cancer genomics data via the Xena platform. Technical report.

[17] Mark D. Robinson, Davis J. McCarthy, and Gordon K. Smyth. edgeR: a Bioconductor package for differential expression analysis of digital gene expression data. Bioinformatics, 26(1):139–140, 1 2010.

[18] Wei Zhang, Shuai Shang, Yingying Yang, Peiyao Lu, Teng Wang, Xinyi Cui, and Xuexi Tang. Identification of DNA methylation-driven genes by integrative analysis of DNA methylation and transcriptome data in pancreatic adenocarcinoma. Exp Ther Med, 19(4):2963–2972, 4 2020.

[19] Philip A. Philip, Ibrahim Azar, Joanne Xiu, Michael J. Hall, Andrew Eugene Hendifar, Emil Lou, Jimmy J. Hwang, Jun Gong, Rebecca Feldman, Michelle Ellis, Phil Stafford, David Spetzler Moh’D M. Khushman, Davendra Sohal, A. Craig Lockhart, Benjamin A. Weinberg, Wafik S. El-Deiry, John Marshall, Anthony F. Shields, and W. Michael Korn. Molecular Characterization of KRAS Wild-type Tumors in Patients with Pancreatic Adenocarcinoma. Clinical Cancer Research, 28(12):2704–2714, 6 2022.

[20] Xiaoyong Fu, Resel Pereira, Carmine De Angelis, Jamunarani Veeraraghavan, Sarmistha Nanda, Lanfang Qin, Maria L Cataldo, Vidyalakshmi Sethunath, Sepideh Mehravaran, Carolina Gutierrez, Gary C Chamness, Qin Feng, Bert W O’Malley, Pier Selenica, Britta Weigelt, Jorge S Reis-Filho, Ofir Cohen, Nikhil Wagle, Agostina Nardone, Rinath Jeselsohn, Myles Brown, Mothaffar F Rimawi, C Kent Osborne, and Rachel Schiff. FOXA1 upregulation promotes enhancer and transcriptional reprogramming in endocrine-resistant breast cancer. Proc Natl Acad Sci USA, 12 2019.

[21] Joanna Obacz, Lucia Sommerova, Daria Sicari, Michal Durech, Tony Avril, Filippo Iuliano, Silvia Pastorekova, Roman Hrstka, Eric Chevet, Frederic Delom, and Delphine Fessart. Extracellular AGR3 regulates breast cancer cells migration via Src signaling. Oncol Lett, 18(5):4449–4456, 11 2019.

[22] Jonathan Ronen, Sikander Hayat, and Altuna Akalin. Evaluation of colorectal cancer subtypes and cell lines using deep learning. Life Science Alliance, 2(6), 12 2019.

